# Endosome positioning coordinates spatially selective GPCR signaling

**DOI:** 10.1101/2022.07.26.501572

**Authors:** Blair K.A. Willette, Nikoleta G. Tsvetanova

## Abstract

G protein-coupled receptors (GPCRs), a class of critical regulators of mammalian physiology, can initiate unique functional responses depending on the subcellular compartment of their activation. Yet, how endosomal receptors transduce location-biased outcomes remains poorly understood. Efforts to uncover the mechanistic basis of compartmentalized GPCR signaling have largely focused on the biochemical aspect of this regulation through dissection of the relevant factors. Here, we assess the biophysical positioning of receptor-containing endosomes as an alternative salient mechanism coordinating the transduction of spatially biased responses. We focus on the prototypical beta2-adrenergic receptor (β2-AR), which preferentially mediates transcriptional reprogramming via cyclic AMP (cAMP) production from early endosomes. We overcome a technical challenge that has hindered the direct assessment of the role of endosome positioning in this paradigm by devising a strategy to selectively and rapidly redistribute endosomes ‘on command’ in intact cells without perturbing their biochemical composition. Next, we present two complementary optical readouts that enable robust measurements of bulk- and gene-specific GPCR/cAMP-dependent transcription with single-cell resolution. We then combine these readouts with rapid endosome relocalization to establish that increasing endosome distance from the nucleus inhibits the initiation of the endosome-dependent response. Lastly, we demonstrate a prominent mechanistic role of phosphodiesterase (PDE)-mediated cAMP hydrolysis in this process. Our study, therefore, illuminates a novel mechanism regulating GPCR function by identifying endosome positioning as a principal mediator of spatially selective receptor signaling.

**Summary:** G protein-coupled receptors (GPCRs) orchestrate essential aspects of mammalian physiology. GPCR function is tightly controlled by endocytic trafficking, where the ligand-activated receptor engages arrestins and clathrin machinery and is subsequently internalized into endosomal compartments^1^. While the endosome-associated receptor pool was classically presumed to be functionally inactive, it is now clear that receptors can also signal from endosomes^2-4^. Moreover, endosomal receptors can initiate cellular responses that are distinct from those activated at the plasma membrane. Transcriptional reprogramming was one of the first location-biased GPCR responses to be identified and shown to be stimulated from intracellular receptors^5, 6^. Since then, compartmentalized signaling has been implicated in the transduction of distinct phosphosignaling^7, 8^ and in the coordination of unique physiologies and drug actions^8-17^. Yet, how the endosome selectively facilitates these responses compared to other subcellular compartments remains unclear.

---

Efforts to mechanistically understand compartmentalized GPCR signaling have focused on the biochemical aspect of this regulation through identification and characterization of relevant factors. Studies have identified regulatory roles for beta and alpha arrestins^16, 18, 19^ as well as trafficking proteins, including sorting nexins and members of the retromer complex^18, 20, 21^, in fine-tuning the activity of endosomal GPCRs. In contrast, the biophysical properties of endosomes that contribute to this phenomenon have been unexplored. Endosomes are highly dynamic organelles located in close proximity to the nucleus. Therefore, their three-dimensional positioning within the cell could play an active role in coordinating GPCR signaling, especially in the context of spatially biased nuclear responses, such as transcriptional signaling. The challenge in testing this model lies in the ability to selectively manipulate the positioning of endosomes without perturbing their biochemical composition.

We address this question by focusing on the beta2-adrenergic receptor (β2-AR), one of the best-characterized GPCRs, which controls cardiovascular and pulmonary physiologies in response to adrenaline and noradrenaline. The β2-AR signals via Gas-dependent stimulation of cyclic AMP (cAMP) from the plasma membrane and endosomes^14^. The endosomal signal preferentially initiates gene transcription through downstream activation and translocation of PKA into the nucleus where it phosphorylates cAMP Response Element-Binding Protein (CREB)^6, 22-24^. Here, we devised a strategy that enables the rapid and selective redistribution of receptor-containing endosomes without disrupting their biochemical integrity to establish the biophysical positioning of endosomes as a primary mediator of location-encoded GPCR signal transduction.

## Results

### A dimerization strategy rapidly redistributes early endosomes toward the cell periphery

To assess the significance of endosome positioning in the activation of spatially biased GPCR signaling, we began by devising a strategy for rapid redistribution of endosomes. Endosome localization is tightly regulated by the cytoskeletal framework of dynein and kinesin motors that move endosomes along microtubules toward the nucleus or cell periphery, respectively^25^. The most commonly used ways to manipulate organelle positioning involve either pharmacologic or genetic perturbations of the cytoskeleton, such as microtubule-depolarizing drugs (e.g., nocodazole), dynein inhibitors (e.g., ciliobrevin D), or depletion/overexpression of motor proteins^26, 27^. However, these approaches are not selectively tailored toward endosomes. In addition, they can lead to cell toxicity and may occur over the course of several days (e.g., in the case of gene depletion/overexpression) and could potentially prompt compensatory genetic mechanisms.

To get around these issues, we set up a chemically inducible dimerization strategy. We fused the early endosome antigen 1 (EEA1) protein to the FKBP-rapamycin-binding (FRB) domain and eGFP to generate EEA1-FRB-eGFP, and the plus-end-directed kinesin motor Kif1A to the FK506 binding protein (FKBP) and tandem dTomato to generate Kif1A-FKBP-tdTomato. In the presence of rapamycin, FRB dimerizes with FKBP, thus bringing together early endosomes and kinesin to facilitate ‘on command’ redistribution of endosomes toward the plasma membrane (**Fig.1A**). For our experiments, we chose HEK293 cells, because they express β2-ARs at native levels^28^ and are an appropriate model to examine receptor signaling under endogenous setting. We transiently transfected HEK293 cells with EEA1-FRB-eGFP and Kif1A-FKBP-tdTomato and used live- and fixed-cell microscopy to evaluate endosome distribution. In comparison to vehicle-treated cells, which exhibited typical endosomal puncta scattered across the cytosol, rapamycin-treated cells showed rapid recruitment of endosomes toward the cell periphery. This redistribution was apparent as early as 5 minutes following bath application of rapamycin and was complete by 30 minutes (**Fig. 1B, Supplementary Video 1**). Quantification of the data revealed a significant rapamycin-dependent increase in the average distance between endosomal puncta and the nucleus (>2-fold increase in mean distance per cell, **Fig. 1C**). In contrast, the dimerization approach did not impact the subcellular distribution of another organelle, the trans-Golgi (**Supplementary Fig. 1**). Thus, our strategy successfully enables selective and rapid redistribution of endosomes in intact cells.

**Figure 1.**
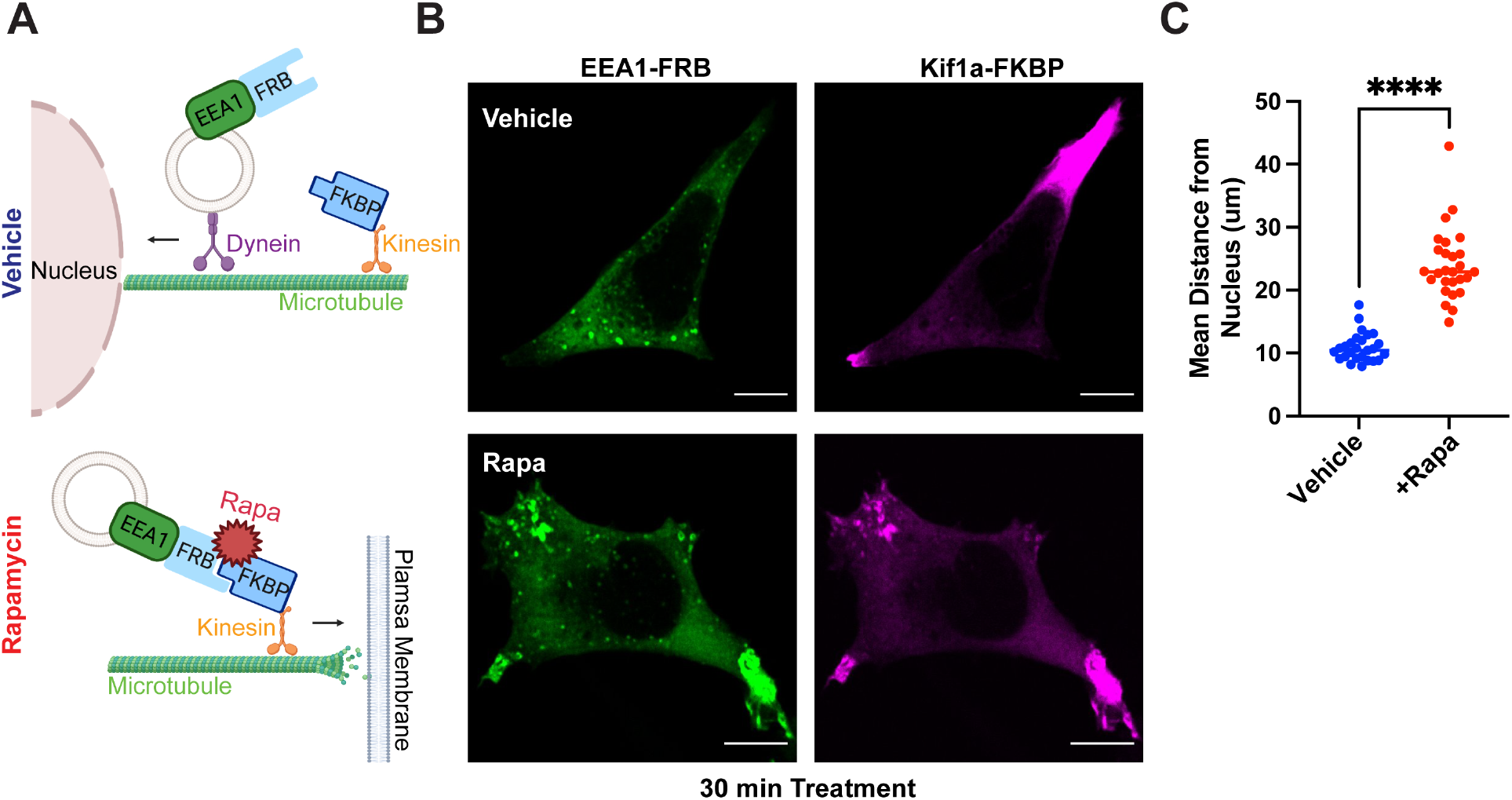
CID approach for redirecting endosomes. **A**. Schematic of the approach. **B-C**. Rapid redistribution of early endosomes by rapamycin treatment. **B**. EEA1-FRB (green), localized at early endosomes, and Kif1a-FKBP (magenta), localized near the cell periphery, visualized by immunofluorescence microscopy in fixed cells. In ethanol-treated cells (vehicle, top), early endosomes are dispersed in the cell. After 1µM rapamycin treatment for 30 min (bottom), early endosomes redistribute to the cell periphery. **C**. Significant increase in mean endosome distance from the nucleus in rapamycin-treated cells. Mean endosome distance per cell = 10.75µm ± 0.46 (EtOH) and 24.04µm ± 1.08 (rapamycin). Data are mean from n = 3 experiments; 27 cells total/condition; **** = *p* < 0.0001 by unpaired Student’s *t-*test. Scale bar = 10µm.

### β2-AR trafficking and signaling are intact in redistributed endosomes

To facilitate the evaluation of endosome positioning in GPCR signaling per se, the dimerization approach should not alter the biochemical composition of endosomes. Hence, we next asked whether the strategy impacted endosomal integrity by assessing β2-AR trafficking and signaling.

To examine β2-AR trafficking, we co-expressed N-terminally Flag-tagged β2-AR and the dimerization constructs, and stimulated receptor internalization using the synthetic agonist isoproterenol. In the absence of agonist, the β2-AR was expressed on the plasma membrane in both vehicle and rapamycin conditions (**Fig. 2A**). In control cells, we observed robust β2-AR endocytosis into EEA1-positive vesicles within 20 minutes of stimulation (**Fig. 2B**, left). The β2-AR internalized similarly in cells pre-treated with rapamycin, despite the dramatic change in endosome positioning (**Fig. 2B**, right and **Fig. 2C**, blue bars). Therefore, rapamycin-induced endosome redistribution does not interfere with proper receptor internalization.

**Figure 2.**
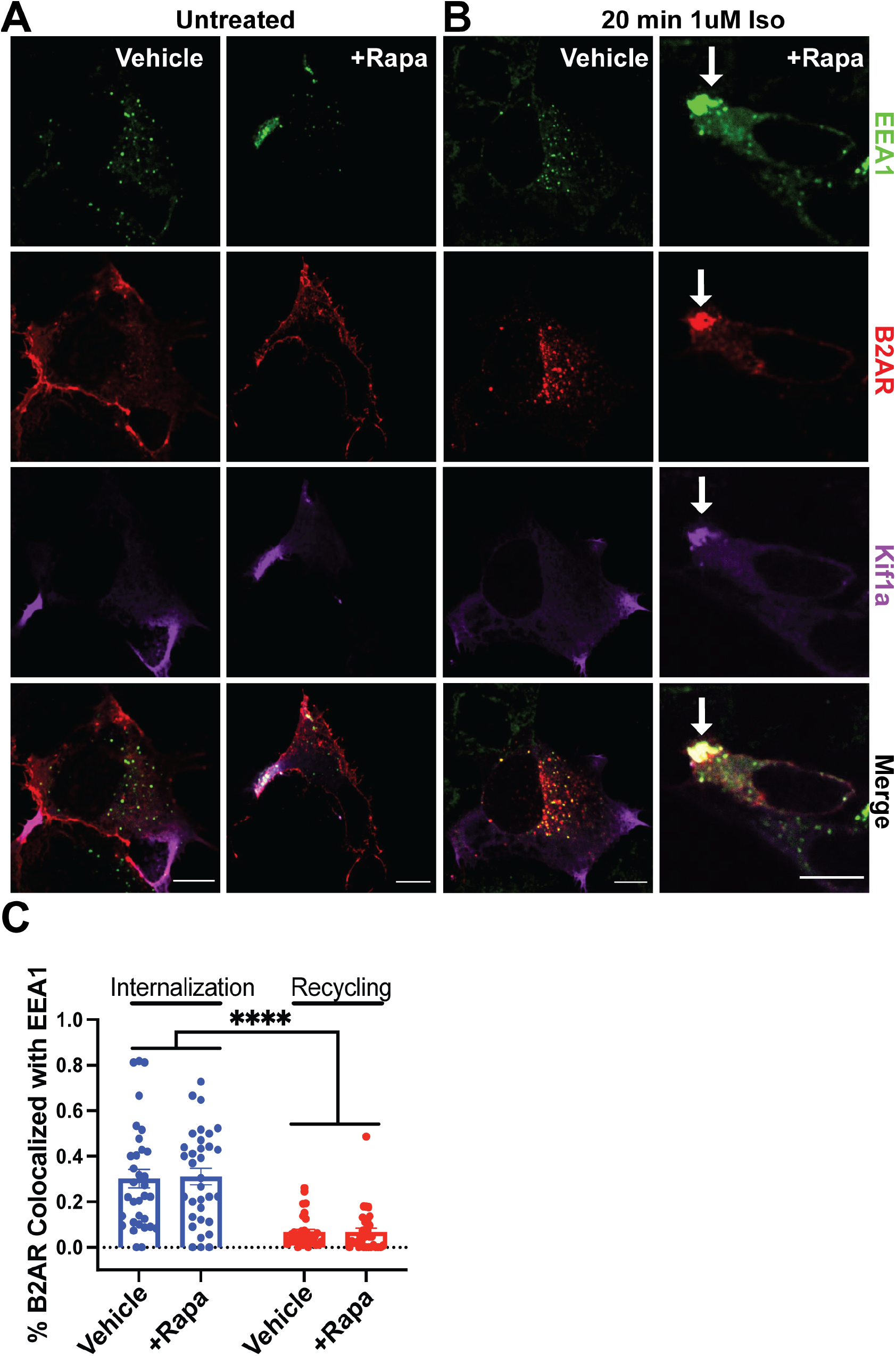
Intact β2-AR trafficking in repositioned endosomes. **A-D**. HEK293 cells were pre-treated with ethanol (“Vehicle”) or 1µM rapamycin (“+Rapa”) for 30 min. Cells were fixed and imaged by immunofluorescence microscopy. **A**. HEK293 cells co-expressing the CID machinery and Flag-β2-AR visualized by immunofluorescence microscopy. β2-AR (red) is present at the cell surface in the absence of β2-AR stimulation. **B**. β2-AR colocalizes with early endosomes (green) after 20 min of activation with 1µM isoproterenol (“Iso”). β2-AR-containing endosomes selectively redistribute toward the cell periphery upon rapamycin application. **C**. Endosome redistribution with rapamycin does not impact β2-AR internalization or recycling compared to vehicle-treated cells. Internalization was induced as describe in **B**. Recycling was induced following internalization by application of 10µM alprenolol for 1 hr (**Supplementary Fig. 2**). Data are mean from n = 3 experiments ± s.e.m.; 33 cells total/condition; **** = *p* < 0.0001 by two-way ANOVA test.

During the initial hours of agonist stimulation, β2-ARs recycle rapidly from early endosomes back to the plasma membrane following endocytosis. This recycling process is mediated by the retromer complex, which is composed of sorting nexins (SNX) and vacuolar protein sorting (VPS) protein^29^. To establish whether β2-AR recycling is impacted by endosome redistribution, we first examined the localization of the endosome-associated retromer component SNX27. SNX27 localized similarly to EEA1-positive endosomal compartments in both control and rapamycin-treated cells (**Supplementary Fig. 2A**). Next, we visualized and quantified recycled β2-ARs in cells stimulated with isoproterenol, then treated with the antagonist alprenolol to induce synchronized recycling (**Supplementary Fig. 2B**). We observed very efficient receptor recycling from redistributed endosomes, which paralleled that seen in normal cells (**Fig. 2C**, red bars). Hence, neither the localization of necessary sorting components nor the recycling of β2-ARs is altered by our approach.

Next, we investigated whether receptors are capable of signaling from redistributed endosomes. We utilized two conformation-selective nanobody biosensors, Nb80 and Nb37, that enable the visualization of active β2-ARs and Gas, respectively^22^. Each nanobody fused to GFP was co-expressed with Flag-β2-AR and the dimerization constructs. In the absence of receptor stimulation, both Nb80 and Nb37 localized diffusively in the cytosol in control and rapamycin-treated cells (**Fig. 3A** and **Supplementary Fig. 3A**). Upon isoproterenol stimulation of control cells, we observed nanobody redistribution to receptor-containing endosomes indicating β2-AR and Gas activation in these compartments (**Fig. 3B**, left and **Supplementary Fig. 3B**, left). We similarly observed co-localization of both biosensors with endosomal β2-ARs in rapamycin pre-treated cells (**Fig. 3B**, right and **Supplementary Fig. 3B**, right). This demonstrates that the β2-AR can adopt an active conformation and stimulate Gas, despite the change in endosome position.

**Figure 3.**
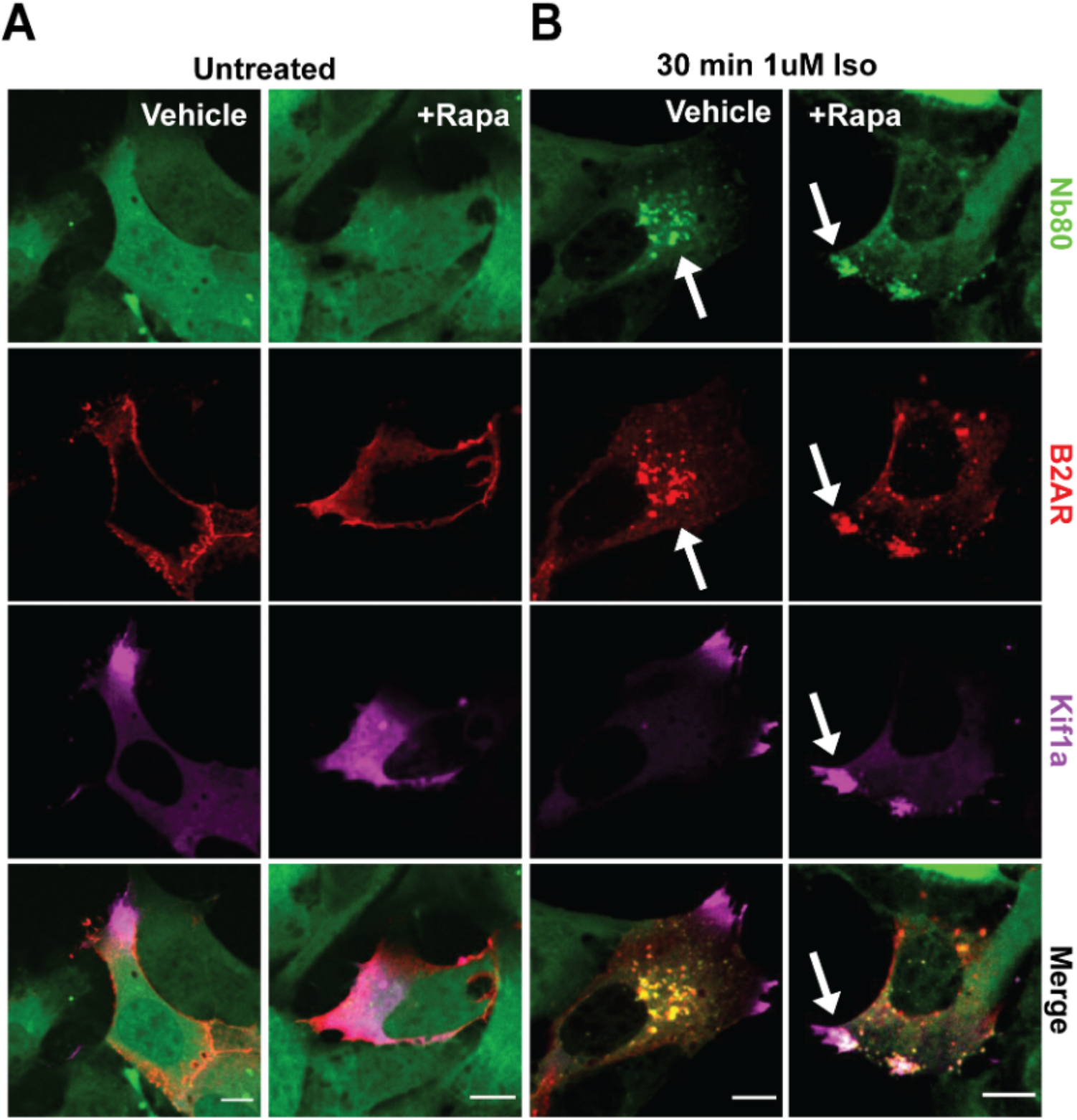
β2-AR adopts an active confirmation on repositioned endosomes. **A-B**. HEK293 cells were pre-treated with ethanol (“Vehicle”) or 1µM rapamycin (“+Rapa”) for 30 min. Flag-β2-AR (red) and Nb80(green) localization in untreated (**A**) and agonist-stimulated CID-expressing cells (**B**). **A**. Nb80 (green) is cytosolic in the absence of β2-AR stimulation. **B**. Redistributed endosomes containing Flag-β2-AR colocalize with Nb80 following 1µM isoproterenol treatment for 30 min. White arrows indicate examples of colocalization between Nb80 with β2-AR. Scale bar = 10µm.

### Microscopy-based readouts report GPCR/cAMP-dependent transcription with single-cell resolution

Endosomal β2-ARs selectively stimulate transcriptional signaling, and we therefore chose to focus on transcription as a pertinent spatially biased GPCR response. Because the dimerization strategy requires optimal co-expression of EEA1-FRB-eGFP and Kif1A-FKBP-tdTomato for efficient endosome relocalization, conventional population-based assays would not be suitable to assess gene transcription. Instead, our experiments necessitated transcriptional readouts compatible with microscopy to allow evaluation of transcriptional signaling in cells that exhibit co-expressed dimerization constructs and efficiently redistributed endosomes.

We began by optimizing an optical readout to monitor the expression of endogenous β2-AR/cAMP target genes. To this end, we utilized proximity ligation in situ hybridization (PLISH), a recently developed approach for single-molecule RNA detection in cells with high sensitivity and specificity^30^. For this analysis, we picked the immediate-early response gene *NR4A1* (Nuclear Receptor Subfamily 4 Group A Member 1), a robust transcriptional target gene of β2-AR activation^6^. We designed four sets of detection probes spanning the length of the transcript and tested them individually and in combination. We achieved the best signal dynamic range when we pooled all four probe sets (isoproterenol vs untreated mean puncta per cell ratio of ∼2.9, **Fig. 4A-B**). Notably, treatment with Dyngo-4a, which inhibits agonist-induced β2-AR internalization (**Supplementary Fig. 4**), drastically blunted the isoproterenol-induced upregulation of *NR4A1* (2.4-fold inhibition, **Fig. 4B**). This is expected for an endosome-dependent response and was previously reported using population-based gene expression readouts^6^. Therefore, single-molecule RNA PLISH analysis of *NR4A1* successfully captures endosomal β2-AR-mediated transcriptional signaling in cells when quantified by microscopy.

**Figure 4.**
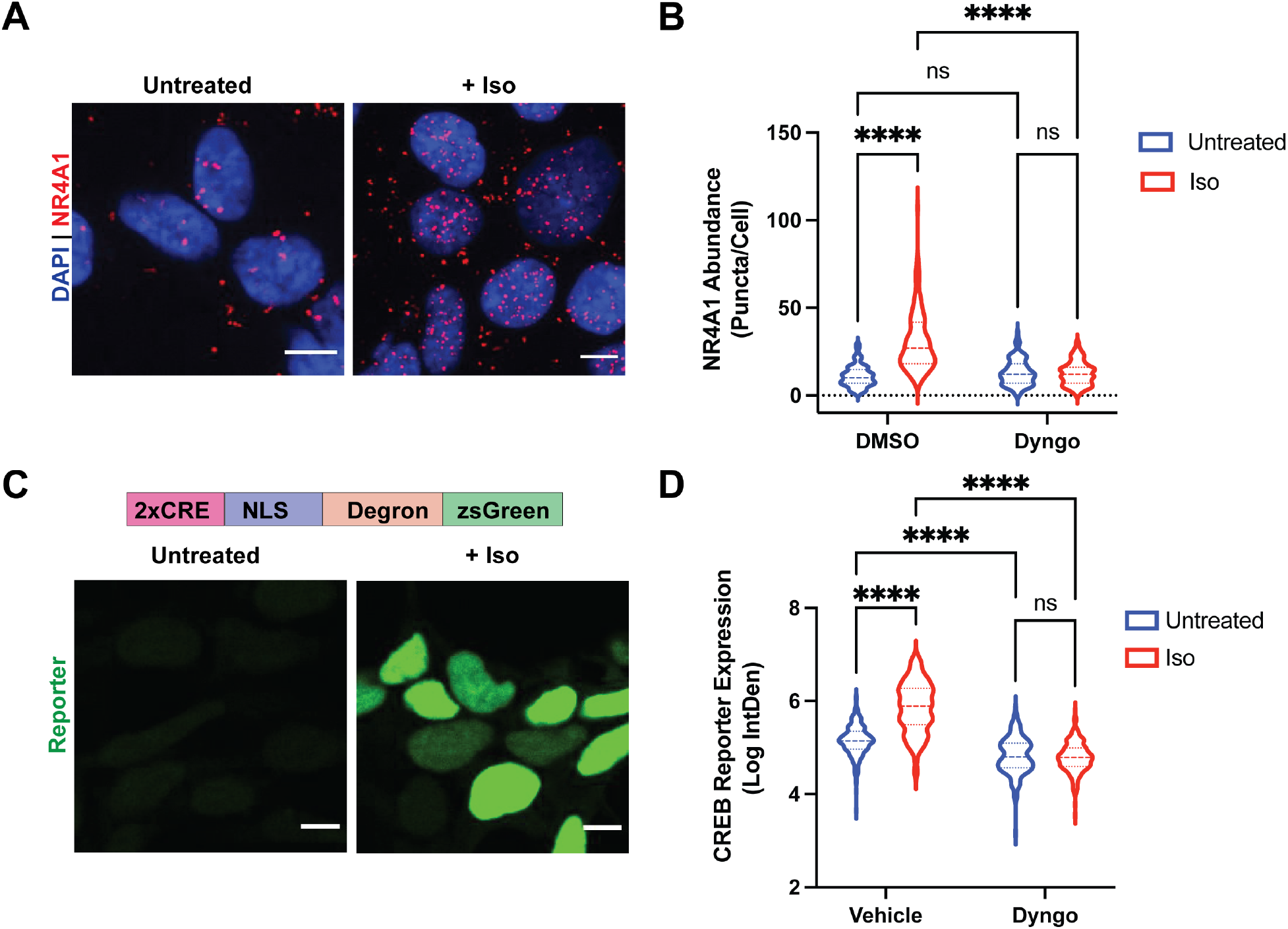
Single-cell optical readouts of GPCR-dependent transcriptional signaling. **A**. Representative images of *NR4A1* mRNA expression by PLISH analysis. Left: untreated; right: treated with 1µM isoproterenol for 2 h. **B**. Quantification of *NR4A1* induction in cells pre-treated with DMSO (Vehicle) or 30µM Dyngo-4A for 20 min, then stimulated with 1µM isoproterenol for 2 h. Data are mean from n = 4 experiments ± s.e.m.; 132 cells total/condition. **C**. Representative images of CREB reporter expression in pCRE-NLS-DD-zsGreen1 cells. Schematic of the reporter is shown on top. CRE = cAMP response element; NLS = nuclear localization sequence. Left: untreated in the presence of 1µM Shield-1 for 4 h; right: treated with 1µM isoproterenol in the presence of 1µM Shield for 4 h. **D**. Quantification of CREB reporter induction in cells pre-treated with DMSO (Vehicle) or 30µM Dyngo-4A for 20 min, then stimulated with 1µM isoproterenol for 4 hr in the presence of 1µM Shield. Data are mean from n = 3 experiments ± s.e.m.; 225 cells total/condition. **** = *p* < 0.0001 by two-way ANOVA test with Tukey. Scale bar = 10µm.

Next, we set off to establish a complementary optical readout of bulk β2-AR/cAMP-dependent transcription. We recently published a fluorescent transcriptional reporter for the cAMP response element-binding protein (CREB)^31^. This reporter contains two cAMP response elements (CREs) recognized by CREB that drive the expression of the zsGreen fluorescent protein and a destabilizing domain (DD). The DD targets the construct for proteasome degradation, but can be selectively stabilized with a cell-permeable compound, Shield-1, thus boosting signal dynamic range^31^. In our previous study, we utilized the reporter in conjunction with flow cytometry. We reasoned that we could apply this readout in combination with microscopy. We reengineered the construct to include a nuclear localization sequence (NLS) for ease of quantification, and generated HEK293 cells stably expressing pCRE-NLS-DD-zsGreen1 by lentiviral transduction. Stimulation with isoproterenol in the presence of stabilizing ligand induced robust reporter accumulation after 4 hours (>4.5-fold relative to vehicle-treated cells in the presence of stabilizing ligand, **Fig. 4C-D**). Similar to the *NR4A1* analysis, this upregulation was abolished by acute blockade of β2-AR internalization with Dyngo-4a (>6-fold inhibition, **Fig. 4D**). Thus, these complementary microscopy-based readouts enable respective measurements of bulk and gene-specific cAMP-dependent transcription with single-cell resolution and can be applied in conjunction with the dimerization system to investigate the role of endosome positioning in spatially biased GPCR signaling.

### Endosome redistribution inhibits GPCR-mediated gene transcription

For all signaling experiments, we utilized the rapalog, AP21967, which permits FRB binding without mTOR cross-reaction^32^ instead of rapamycin. Notably, pretreatment of cells with rapalog did not impact basal levels of transcription compared to control-treated cells (log-transformed basal levels of 6.9 vs. 7.2 and 4.94 vs. 4.87 for rapalog-vs control-treated cells in PLISH and CREB reporter assays, respectively; **Fig. 5A and 5B**). Therefore, rapalog treatment alone does not affect basal *NR4A1* levels measured by RNA in-situ hybridization and CREB signaling measured with the fluorescent reporter.

**Figure 5.**
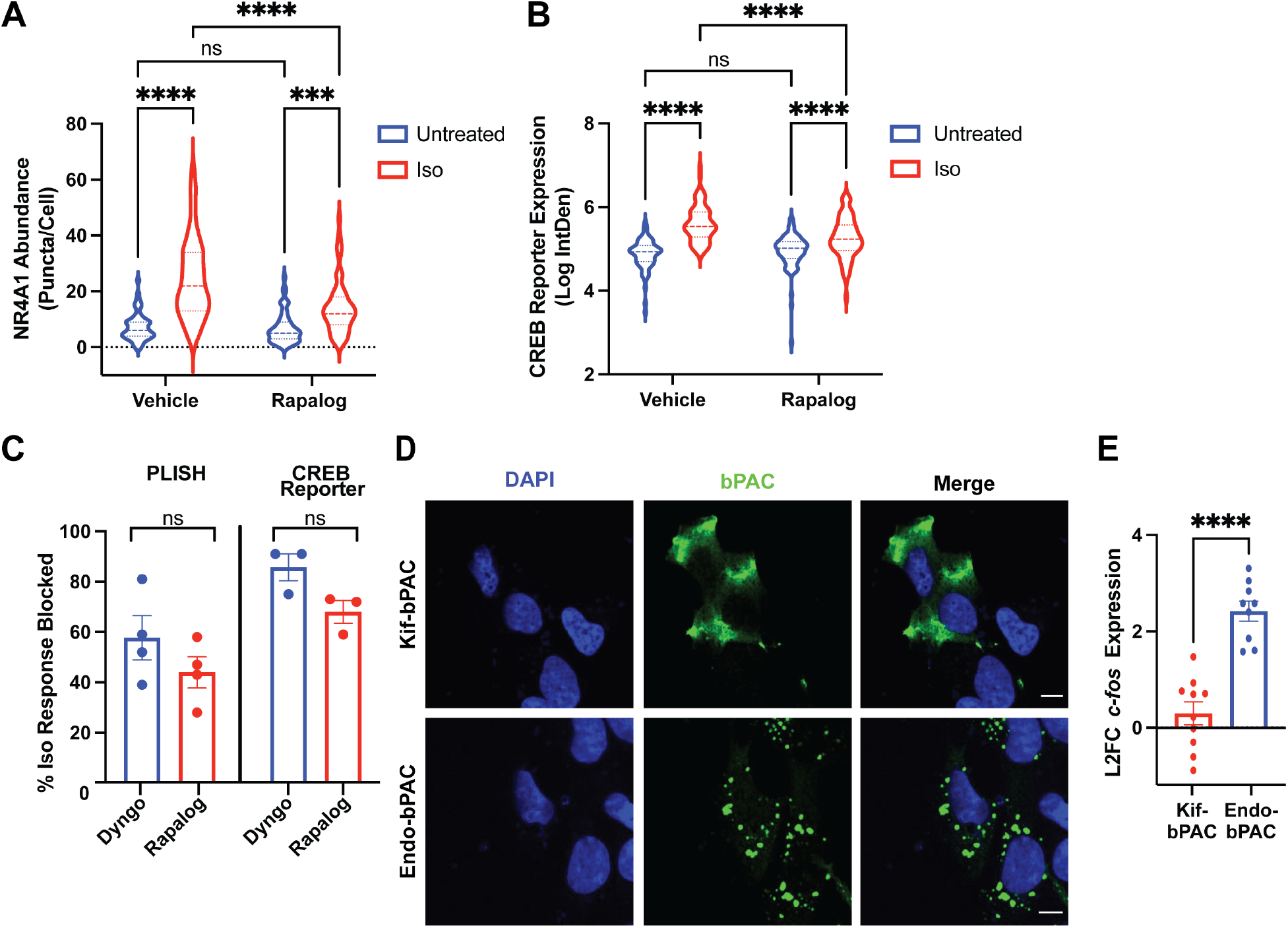
Endosome redistribution blunts site-selective GPCR signaling. **A-C**. Cells expressing the CID system were pre-treated with ethanol (“Vehicle”) or 1µM AP21967 (“Rapalog”) for 30 min before agonist stimulation. **** = *p* < 0.0001 by two-way ANOVA test with Tukey. **A**. *NR4A1* mRNA expression by PLISH analysis of untreated or isoproterenol-stimulated cells (“Iso”, 1µM, 2 h). Data are mean from n = 4 experiments ± s.e.m.; 55 cells total/condition. **B**. CREB reporter expression in pCRE-NLS-DD-zsGreen1 cells untreated in the presence of 1µM Shield or stimulated with 1µM isoproterenol (“Iso”) in the presence of 1µM Shield for 4 h. Data are mean from n = 4 experiments ± s.e.m.; 70-77 cells total/condition. **C**. Endosome positioning regulates the initiation of cAMP-dependent gene transcription to a comparable extent as β2-AR endocytosis. The % inhibition of Isoproterenol-induced response for PLISH (left) and CREB reporter (right) assays. Data are mean from n = 3-4 experiments ± s.e.m. and analyzed by unpaired Student’s *t-*test. **D**. Representative images of bPACs (green) localized to the periphery by fusion to a kinesin motor (“Kif-bPAC”, top) or to the early endosome by fusion to a 2X FYVE domain (“Endo-bPAC”, bottom). Cells were visualized by immunofluorescence microscopy following staining with Alexa 647-conjugated anti-myc antibody. **E**. *c-Fos* induction measured by RT-qPCR analysis after bPAC photostimulation for 3 min. Data represent mean from n = 9-10 experiments ± s.e.m. **** = *p* < 0.0001 by unpaired Student’s *t-*test. Scale bar = 10µm.

Next, we analyzed the impact of endosome relocalization on the endogenous receptor-induced transcriptional responses using PLISH. In control cells transfected with the two dimerization constructs, we observed robust isoproterenol-dependent *NR4A1* accumulation (3.4-fold, **Fig. 5A, Supplementary Fig. 5A**, left). Remarkably, endosome redistribution significantly blunted this response (1.7-fold decrease, **Fig. 5A** and **Supplementary Fig. 5A**, right). At the same time, the expression of the housekeeping gene *GAPDH*, which is not regulated via the β2-AR cascade, was not impacted by the manipulation (**Supplementary Fig. 5B**).

Then, we examined transcriptional signaling in the pCRE-NLS-DD-zsGreen1 cell line. In agreement with the results from the PLISH analysis, here we saw greater than 5-fold induction of CREB reporter accumulation following isoproterenol treatment in control cells expressing the dimerization system, which in turn was dampened by pre-application of rapalog (>2-fold decrease, **Fig. 5B** and **Supplementary Fig. 5C**). On the other hand, we observed comparable reporter induction following direct stimulation of CREB using the cell-permeable cAMP analog, 8-CPT-cAMP, which bypasses the need for receptor and adenylyl cyclase activation (**Supplementary Fig. 5D**). Therefore, the inhibition is specific for the GPCR pathway.

Based on these outcomes, we next quantified the extent to which endosome positioning contributes to the spatial bias of GPCR signaling. Since Dyngo-4a severely blocked endocytosis of β2-ARs (**Supplementary Fig. 4**), we defined % loss of isoproterenol-induced signaling in Dyngo-4a-treated cells to represent the endosome-dependent aspect of the transcriptional response. We observed ∼60% and ∼85% of endosome-dependent transcriptional signaling measured using PLISH and the CREB reporter, respectively (**Fig. 5C**, blue bars). Strikingly, redistribution of β2-AR-containing endosomes mirrored very closely the extent of inhibition observed with Dyngo-4a (∼40% and ∼70% by PLISH and CREB reporter analysis, respectively; **Fig. 5C**, red bars). Hence, endosome positioning is the major determinant of endocytosis-dependent transcriptional signaling in response to GPCR stimulation.

### The subcellular position of cAMP accumulation coordinates activation of transcriptional signaling

As an orthogonal approach to investigate the significance of the biophysical location of signaling, we employed a previously described optogenetic strategy for localized cAMP production^6, 7^. In this approach, the bacteria-derived photoactivatable adenylyl cyclase, bPAC, is fused to localization-specific targeting sequences to enable spatiotemporal control of second messenger accumulation. We reasoned that bPAC fused to the FYVE domain derived from Hrs^33^ (“Endo-bPAC”) would mimic signals originating from natively localized endosomes, while bPAC fused to a Kif1A-derived targeting sequence (“Kif-bPAC”) would mirror signals from the subcellular site of repositioned endosome accumulation. We confirmed proper construct expression and localization (**Fig. 5D, Supplementary Fig. 6A-C**). Then, we measured cAMP concentrations downstream of Kif-bPAC and Endo-bPAC with a colorimetric immunoassay. This analysis showed that comparable cAMP was generated from each site following photostimulation with matched light doses (**Supplementary Fig. 6D**). Next, we examined transcriptional signaling by quantitative PCR analysis of an established β2-AR target gene, *c-fos*^*6*^. Photostimulation of Endo-bPAC activated robust signaling, while Kif-bPAC activated a very weak transcriptional response (**Fig. 6E**). More importantly, this trend held across a range of light doses (**Supplementary Fig. 6D-E**). These results are consistent with the outcome from our analysis of endosomal positioning and suggest that cAMP signals originating at a greater distance from the nucleus relative to natively localized endosomes are effectively uncoupled from downstream transcriptional control.

**Figure 6.**
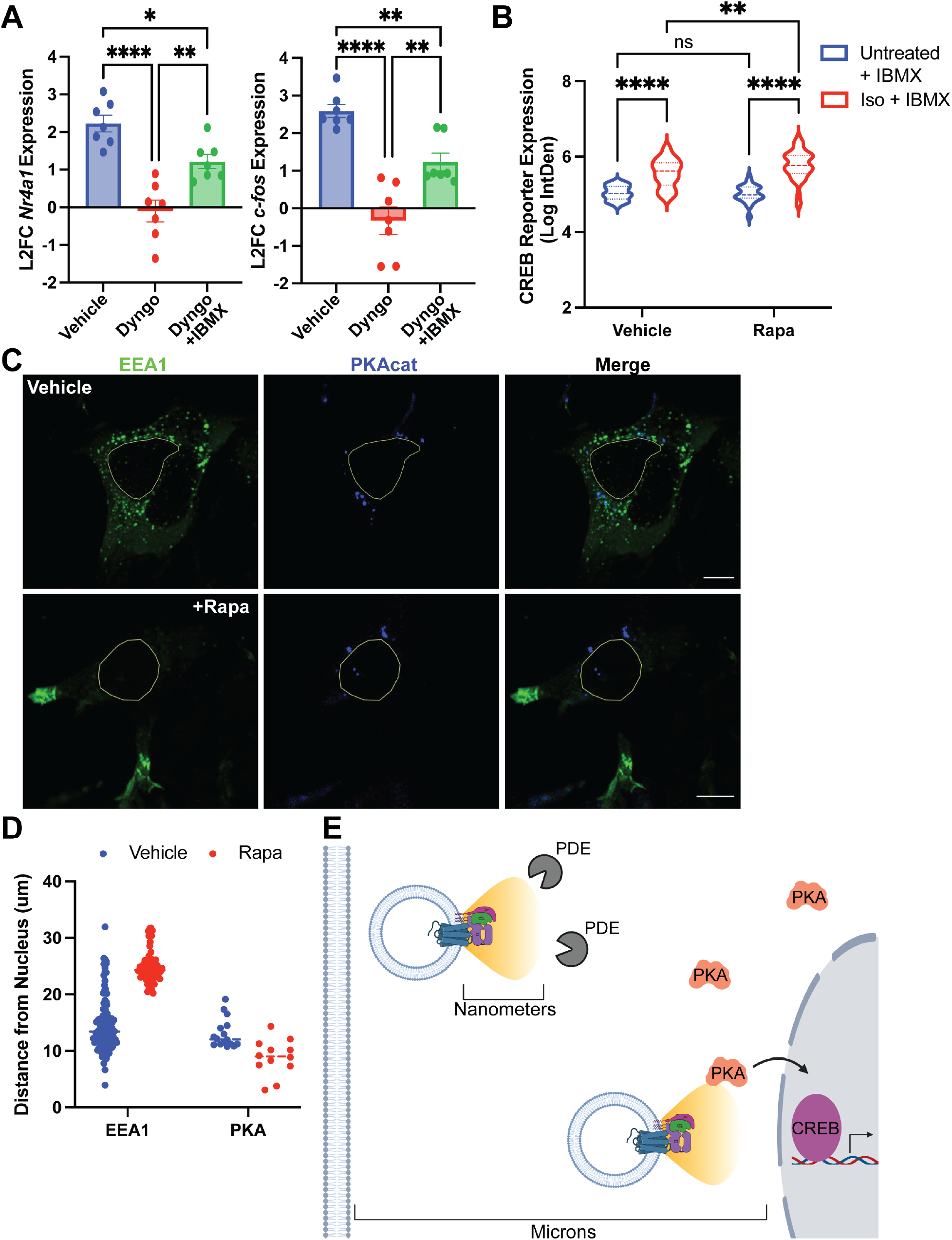
Endosomes precisely position receptors with respect to relevant signaling effectors. **A**. *c-Fos* and *NR4A1* induction measured by RT-qPCR analysis after β2-AR activation with 1µM isoproterenol for 1 h. Cells were pretreated with DMSO (vehicle) or 30µM Dyngo-4A for 20 min, then stimulated in the absence or presence of 100µM IBMX. Data represent mean from n = 7 experiments ± s.e.m. **** = *p* < 0.0001, ** = *p* < 0.01, * = *p* < 0.05 by one-way ANOVA test with Tukey. **B**. CREB reporter expression in pCRE-NLS-DD-zsGreen1 cells stimulated with 1µM isoproterenol in the presence of 1µM Shield for 4 hr with or without 100µM IBMX. Data are mean from n = 3 experiments ± s.e.m.; 45 cells total/condition. **** = *p* < 0.0001, ** = *p* < 0.01 by two-way ANOVA test with Tukey. **C**. Representative images of PKA_cat_-BFP (blue) and EEA1-FRB (green) visualized by immunofluorescence microscopy in fixed cells treated with ethanol (“Vehicle”, right) or 1µM rapamycin (“+Rapa”, left) for 20 min. **D**. Quantification of the distance of EEA1 clusters (left) and PKA particles (right) from the nucleus in response to ethanol (“Vehicle”, blue) or 1µM rapamycin (“+Rapa”, red) based on the representative image shown in **C. E**. Model: Endosomes enable site-selective outputs by functioning as vehicles to deliver the receptor in proximity to PKA and the nucleus and away from phosphodiesterases. Scale bar = 10µm.

### PDE activity buffers GPCR-mediated gene transcription from repositioned endosomes

To determine how endosome localization coordinates receptor signaling, we interrogated the role of phosphodiesterases (PDEs), a well-established class of GPCR/cAMP signaling effectors. PDEs hydrolyze cyclic nucleotides and therefore play a critical role in signal compartmentalization^34^. Therefore, we hypothesized that PDE hydrolysis may contribute to the spatial bias of GPCR-driven transcription and the regulatory role endosome positioning plays in the process. To test this model, we investigated transcriptional activation under two scenarios. First, we confined the β2-AR to the plasma membrane with Dyngo-4a and measure transcriptional signaling by RT-qPCR analysis. As previously reported, endocytic blockade inhibited gene upregulation downstream of activated β2-ARs (**Fig. 6A**)^6^. However, this inhibition was successfully reversed by co-application of the broad-spectrum phosphodiesterase inhibitor, 3-isobutyl-1-methylxanthine (IBMX) (**Fig. 6A**). In parallel experiments, we tested the impact of PDE inhibition on CREB reporter expression in cells with natively localized versus repositioned endosomes. We observed that IBMX similarly eliminated the rapalog-dependent differences in CREB reporter accumulation in pCRE-NLS-DD-zsGreen1 cells treated with isoproterenol (**Fig. 6B**). Therefore, PDE-dependent cAMP hydrolysis prevents the propagation of receptor signals originating beyond the sites of natively localized endosomes.

## Discussion

Despite an emergent recognition of the functional role of endosomal GPCRs, how intracellular receptors transduce distinct responses has remained an open question. In principle, endosomes could selectively enable transcription by presenting a unique biochemical environment or due to their three-dimensional localization, which provides proximity to the nucleus and to relevant signaling effectors. In the present study, we devised an approach to distinguish between these models. A rapamycin-dependent dimerization strategy dramatically redistributed endosomes away from the nucleus without perturbing the complexes required for proper receptor trafficking and signaling, including the localization of an early endosome-resident sorting protein (**Fig. 1, 2 and 3, Supplementary Fig. 2 and 3**). Coupled with two complementary approaches to measure β2-AR signaling with single-cell resolution and high reproducibility, this system allowed us to rigorously dissect the role of endosome positioning in receptor-mediated gene transcription.

Rapid endosome redistribution drastically blunted transcriptional signaling downstream of the β2-AR (**Fig. 5**). Strikingly, the magnitude of inhibition closely mirrored the phenotype observed upon complete blockade of receptor internalization (∼45% and 70% inhibition upon endosome relocalization versus ∼55% and 85% with Dyngo-4a using PLISH or the fluorescent transcriptional readout, respectively; **Fig. 5C**). In addition, the inhibition was selective for the receptor cascade, because it could be circumvented by direct stimulation with a cAMP analog (**Supplementary Fig. 5D**). Therefore, our data argue that the biophysical localization of endosomes per se is a primary mechanism underlying the ability of these organelles to propagate spatially biased GPCR responses. We note that Dyngo-4a precludes not only β2-AR endocytosis but also subsequent receptor resensitization via recycling. In contrast, β2-ARs recycle efficiently out of repositioned endosomes (**Fig. 2**). Therefore, our analysis (**Fig. 5C**) may in fact underestimate the contribution of endosome positioning to transcriptional signaling. Our findings, in turn, fit a broader appreciation for the significance of organelle positioning in cellular physiology. For example, lysosome positioning was found to regulate mTORC1 signaling to subsequently impact autophagosome formation^26^. Similarly, retrograde endosome movement was required for neurotrophic factor signal transmission from distal axons to the nucleus to stimulate various responses, including CREB-dependent transcription^35^.

We propose that the positioning of endosomal receptors plays an essential regulatory role because of the highly compartmentalized nature of the cAMP cascade, which limits the initiation of downstream responses. PKA signaling is reported to be restricted to nano-sized ‘hotspots’^36^. This compartmentalization happens through a number of mechanisms. First, PKA is ‘trapped’ on organelle membranes via the actions of A-kinase anchoring proteins^37^ and in phase-separated biomolecular condensates^38^. Further, signaling is buffered by PDEs, which efficiently hydrolyze cAMP within a narrow radius^34, 39^. In addition, PKA requires a high in-cell cAMP activation threshold, recently reported to be in the micromolar range^40^, which renders local PDE actions especially relevant. Collectively, these properties of the cascade allow the initiation of cAMP- and PKA-dependent responses to be spatially controlled. Based on this and our data, we propose a model according to which endosomes provide the optimal location for the propagation of the GPCR signal in two ways. First, they allow the receptor/signaling complex to evade PDE hydrolysis, which we recently showed to be higher at the cell periphery relative to where endosomes are natively located^7^. Indeed, inhibition of phosphodiesterase activity rescued the inability of receptor signals originating from relocalized endosomes to activate downstream transcription (**Fig. 6B**). Second, endosomes position the receptor in direct proximity to the nucleus (**Fig. 1C**) and to the relevant downstream effector PKA. In agreement with previous reports^10, 24, 38^, we observe the catalytic subunit of PKA, PKA_cat_, on intracellular puncta proximal to natively localized endosomes (**Fig. 6C-D**). The localization of PKA_cat_ puncta does not change upon endosome redistribution (**Fig. 6C-D**). Hence, cAMP produced via the receptor/Gas at repositioned endosomes is simultaneously buffered by active PDE hydrolysis while also having to propagate longer distances to reach its spatially confined downstream effector. We propose that these events ultimately synergize to preclude the initiation of transcriptional signaling (**Fig. 6E**).

The experimental platform described here could be applied in future analyses to determine how endosome positioning impacts other important outputs of the GPCR cascade. Both G protein-dependent and G protein-independent responses can be initiated from endosomal compartments^2, 41^. Based on the model proposed above, we speculate that additional G protein/cAMP-stimulated responses would also be preferentially stimulated by endosomal GPCR activation and coordinated by endosome positioning. Indeed, it was recently shown that phosphosignaling is driven primarily by endosomal cAMP and requires receptor internalization^7^. However, it was not tested whether these processes are dependent on the precise localization of receptor-containing endosomes. Further, it would be particularly interesting to examine whether endosome positioning plays a role in arrestin-dependent signaling (e.g., arrestin stimulation of ERK1/2 signaling^42^), which is not orchestrated through the actions of PKA, cAMP and PDEs. The two complementary approaches to measure GPCR-dependent transcription by microscopy would also be valuable to further dissect the regulatory logic of the receptor cascade. These functional readouts can be applied to interrogate factors important for endosome positioning or receptor trafficking in and out of endosomes. Because of the single-cell nature of the readouts, we anticipate that these would also offer the unprecedented ability to examine pathway regulation in parallel across different cell types in a heterogeneous setting.

In summary, our study identifies a novel mechanism underlying spatially biased GPCR signaling by demonstrating an essential role for endosome positioning in the propagation of site-selective transcriptional signaling. Our findings were corroborated by two complementary approaches, including specific readouts for β2-AR activity and a receptor-independent optogenetic method for cAMP generation. Therefore, these results likely represent a principle of broader significance that would apply to an ever-growing list of Gas/cAMP-coupled GPCRs found to signal from endosomes^9, 16, 43-45^.

## Materials and Methods

### Chemicals

(-)-Isoproterenol hydrochloride was purchased from Sigma-Aldrich (Cat #I6504), dissolved in water/100mM ascorbic acid to 10mM stock, and used at 1µM final concentration. Alprenolol hydrochloride was purchased from Sigma-Aldrich (Cat # A0360000), dissolved in DMSO to 10mM stock, and used at 10µM final concentration. DD stabilizing compound, Shield-1, purchased from Takara Bio (Cat #632189) and used at 1µM final concentration. 8-(4-Chlorophenylthio)adenosine 3’,5’-cyclic monophosphate (8-CPT-cAMP) was purchased from Abcam (Cat #ab120424), dissolved in water to 150mM stock, and used at 150µM final concentration. 3-isobutyl-1-methylxanthine (IBMX) was purchased from Sigma (Cat #I5879-1G), dissolved in DMSO to 100mM stock, and used at 100µM final concentration. Rapamycin was purchased from Sigma-Aldrich (Cat #553210-1MG), dissolved in ethanol to 10mM stock, and used at 1µM final concentration. Rapalog (A/C Heterodimerizer) was purchased from Takara (Cat #635056) and used at 1µM final concentration. Dyngo-4a was purchased from Abcam (Cat #ab120689), dissolved in DMSO to 30mM stock, and used at 30µM final concentration in serum-free medium.

### Construct cloning

A previously described lentiviral plasmid encoding a transcriptional reporter for CREB activity^31^ was used as the backbone for the CREB transcriptional reporter in this study. To express this construct in the nucleus, a nuclear localization sequence (NLS) was amplified from dCas9-BFP-KRAB (a gift from Martin Kampmann, UCSF) and inserted into the BamHI-digested reporter backbone using Gibson cloning. pBa-Kif1A(1-396)-tdTomato-FKBP was generated by Asc1/Hpa1 restriction digest of pBa.Kif1a 1-396.GFP (backbone; Addgene, Cat #45058) and of pBa-KIF5C 559-tdTomato-FKBP (insert; Addgene, Cat #64211) followed by ligation. CMV-EGFP-FRB-EEA1 was generated by site-directed mutagenesis to remove the stop codon upstream of FRB from pEGFP-FRB (Addgene, Cat #25919). Next, EEA1 was amplified from GFP-EEA1 (Addgene, Cat #42307) and Gibson cloning was used to insert the product into the BsiWI-digested pEGFP-FRB backbone. FRB-EEA1 was generated by site-directed mutagenesis to insert an AgeI cut site upstream of EGFP in CMV-EGFP-FRB-EEA1. AgeI digestion was used to remove EGFP, followed by ligation into the CMV-FRB-EEA1 backbone. CMV-Kif1A-bPAC was generated by PCR amplification of Kif1A(1-396) and Gibson cloning into the Xbal/EcoRV-digested CMV-bPAC backbone described previously^6^. To clone Nb37 into a lentivirus backbone, Nb37-GFP^22^ was amplified and inserted into the NheI/XbaI-digested pMK1200 backbone (a gift from Martin Kampmann, UCSF) by Gibson cloning. Similarly, Nb80-GFP^22^ was amplified and inserted into the NheI/XbaI-digested pMK1200 backbone by Gibson cloning to generate Nb80 for lentivirus production.

### Cell culture and transcriptional reporter cell line generation

HEK293 and HEK293T cells were obtained from ATCC and grown at 37°C/5% CO_2_ in Dulbecco’s Modified Eagle Medium (4.5 g/L glucose and L-glutamine, no sodium pyruvate; Thermo Fisher Scientific Cat # 11965118) and supplemented with 10% fetal bovine serum. HEK293 cells stably expressing pCRE-NLS-DD-zsGreen1 were generated by lentiviral transduction and fluorescence activated cell sorting for zsGreen1+ in the 488 channel on a BD DIVA (BD Biosciences). Ten GFP-positive cells were sorted per well in a 96-well plate and expanded. Populations were then tested by flow cytometry (BD FACSCanto II) for cAMP-induced GFP expression following stimulation with 1µM isoproterenol in media containing 1µM Shield-1 for 4 hr. We selected the population with the highest signal dynamic range for our follow-up experiments.

### Lentivirus production

Lentivirus was used to generate the pCRE-NLS-DD-zsGreen1 cell line and to express Nb37, Nb80, and Flag-β2-AR. To make lentivirus, HEK293T cells were transfected with pCRE-NLS-DD-zsGreen1, EF1-Nb37-GFP, EF1-Nb80-GFP, or pFUGW-Flag-β2-AR (a gift from Mark von Zastrow, UCSF) lentivirus vectors and standard packaging vectors (VSVG and psPAX2) using Lipofectamine 2000 (ThermoFisher, Cat #1668027) following recommended protocols. Virus-containing supernatants were collected 72 hr after transfection, filtered through a 0.45µ SFCA filter, and either used on the same day or concentrated using Lenti-X concentrator solution (Takara Bio, Cat #631231) and snap frozen at −80C.

### Spinning disk confocal imaging

Live- and fixed-cell imaging were performed on Andor Dragonfly Spinning Disk Confocal microscope with 405 nm, 488 nm, 561 nm, 637 nm diode lasers (Andor). The microscope was fitted with 40x/1.3 HC PL APO CS2 (Leica, Cat #11506358), Oil, WD: 0.24mm; 63x/1.47 TIRF HC PL APO CORR (Leica, Cat # 11506319), Oil, WD: 0.10mm; and 100x/1.40-0.70 HCX PL

APO (Leica, Cat # 11506210), Oil, WD: 0.09mm objectives. The Andor iXon Life 888 1024×1024 EMCCD, pixel size: 13µm camera was used with and without 2X zoom optics. Live cell imaging took place in a 37°C and 5% CO_2_ chamber (Okolab).

### Early endosome redistribution assay and quantification

Cells plated on Poly-L-lysine (Sigma, Cat #P8920-100ML) coated coverslips (Thomas Scientific, Cat #217N81) were transfected with Kif1A-tdTomato and EGFP-FRB-EEA1 using Lipofectamine 2000 (ThermoFisher, Cat #11668027). The following day, the cells were treated with rapamycin or ethanol vehicle for 20 min. Following treatment, cells were fixed using 3.7% formaldehyde (ThermoFisher, Cat #28908) in modified BRB80 buffer (80mM PIPES pH 6.8, 1mM MgCl_2_, 1mM CaCl_2_) for 20 min. Fixation was quenched and cells were permeabilized using 2.5%milk/0.1% Triton X-100 (Sigma, Cat #×100-100ML) in TBS for 20 min. Primary antibodies [Anti-EEA1 (Cell Signaling, Cat #3288S), Anti-SNX27 (Abcam, Cat #ab77799), Anti-TGN46 (Abcam, Cat #ab50595)] were diluted 1:500 in permeabilization solution and incubated for 1 hr at room temperature. Secondary antibodies [Donkey anti-Mouse IgG (H+L) Secondary Antibody, Alexa Fluor 647 (Invitrogen, Cat #A-315710); Donkey anti-Rabbit IgG (H+L) Secondary Antibody, Alexa Fluor 647 (Invitrogen, Cat #A-31573)] were diluted 1:1000 in permeabilization solution and incubated for 30 min at room temperature, protected from light. Following two washes in permeabilization solution and one wash in PBS, coverslips were mounted using ProlongGold with DAPI (Thermo Scientific, Cat #P36931). Slides were imaged on an Andor Dragonfly Spinning Disk Confocal microscope. Images were imported into FIJI. Single cells were isolated by drawing ROIs around the cell periphery seen in the Kif1A-containing channel and nuclei were identified by drawing ROIs of DAPI-stained nuclei. Clusters of EEA1-containing endosomes were segmented using the Morphological Segmentation plugin (MorphoLibJ)^46^. Using the distance formula, the distance from segmented EEA1 particles to the nucleus center of mass was calculated based on particle and nuclear x,y coordinates. Distances of all endosomes to nucleus were averaged per cell.

### β2-AR trafficking and quantification

Trafficking was quantified by two methods. For microscopy-based analysis, HEK293 cells plated on Poly-L-lysine-coated coverslips were transfected with Kif1A-tdTomato-FKBP, EGFP-FRB-EEA1, and Flag-β2-AR using Lipofectamine 2000. Cells were first treated with either rapamycin or vehicle (ethanol) for 20 min to induce redistribution, and then supplemented with Alexa 647-conjugated M1 antibody (1:1000) for 10 min to label cell-surface Flag-β2-ARs. To reduce non-specific antibody binding, the media was changed with fresh pre-equilibrated medium containing rapamycin or ethanol and either isoproterenol or vehicle for 20 min to induce β2-AR internalization. Following receptor internalization, media was changed with rapamycin or ethanol control with or without alprenolol for 1 hr to induce β2-AR recycling. Cells were fixed as described above and slides were imaged on Andor Dragonfly Spinning Disk Confocal microscope. Images were imported into FIJI and individual channels were saved as TIFF files. The Squassh plugin (MosaicSuite)^47^ was used to remove background, apply a joint deconvolution-segmentation procedure, and calculate object-based colocalization using information about the shapes and intensities of all objects in both channels. From the supplied R script, the C(number) was used to estimate internalized or recycled β2-ARs as the ratio of (number of β2-AR puncta colocalized with EEA1)/(total EEA1 puncta).

For flow cytometry-based analysis, we generated HEK293 cells stably expressing Flag-β2-AR and treated with 30µM Dyngo-4a or DMSO in serum-free DMEM for 20 minutes. Next, cells were treated either isoproterenol or vehicle for 30 min to induce β2-AR internalization. Cells were washed with PBS and then incubated with Alexa 647-conjugated M1 antibody (1:1000) for 1 hr at 4°C, shaking at 200 rpm. Then, cells were lifted, placed in tubes for flow cytometry (BD FACS Canto2), and 10,000 cells were analyzed per sample. The gated Alexa 647 mean of the singlet population was used as the amount of receptor surface expression. The % internalized receptors was calculated by 100%-(# surface receptors after 30 min isoproterenol)/(# surface receptors after vehicle) × 100%.

### Nanobody Colocalization in Relocalized Endosomes

HEK293 cells were infected with either EF1-Nb80-GFP or EF1-Nb37-GFP. Three days later, cells were plated on Poly-L-lysine-coated coverslips. The following day cells were transfected with Kif1A-tdTomato-FKBP, EGFP-FRB-EEA1, HA-Gas (Nb37 expressing cells only), and Flag-β2-AR using Lipofectamine 2000. Cells were first treated with either rapamycin or vehicle (ethanol) for 30 min to induce redistribution, and then supplemented with Alexa 647-conjugated M1 antibody (1:1000) for 10 min to label cell-surface Flag-β2-ARs. To reduce non-specific antibody binding, the media was changed with fresh pre-equilibrated medium containing rapamycin or ethanol and either isoproterenol or vehicle for 30 min to induce β2-AR internalization. Cells were fixed as described above and slides were imaged on Andor Dragonfly Spinning Disk Confocal microscope. Images were imported into FIJI and individual channels were saved as TIFF files. Since these experiments necessitated the transfection of multiple constructs, we observed reduced expression of the dimerization components and, consequently, decreased efficiency of endosome redistribution in cells pre-treated with rapamycin. Nevertheless, a sufficient fraction of endosomes relocated to the cell edge in rapamycin-dependent manner to allow evaluation of Nb80 and Nb37 recruitment and demonstration of functional β2-AR signaling.

### NR4A1 expression PLISH analysis

HEK293 cells were plated on Poly-L-lysine-coated coverslips and transfected with Kif1A-tdTomato-FKBP and EGFP-FRB-EEA1 using Lipofectamine 2000. When inhibiting receptor internalization, cells were first treated with 30µM Dyngo-4a or DMSO in serum-free DMEM for 20 min. When relocalizing endosomes, cells were first treated with either rapalog or ethanol control in serum-free DMEM media for 20 min. Next, cells were treated with isoproterenol or vehicle control for 2 hr. Cells were fixed using 3.7% formaldehyde in PBS/0.1%DEPC (Sigma-Aldrich, Cat #D5758-5ML) for 20 min. Coverslips were washed three times with PBS, dehydrated by an ethanol dilution series, air dried for 10 min, and enclosed by application of a seal chamber (Grace Bio Labs, Cat #621505) as previously described^30^.

The following H-probe oligonucleotides were used for *NR4A1:*

HL4X-NR4A1-1879 (TTAGTAGGCGAACTTACGTCGTTATGTTGTCAATGATGGGTGGAGG), HR4X-NR4A1-1899 (GCAGCGTGTCCATGAAGATCTTATACGTCGAGTTGAACATAAGTGCG), HL4X-NR4A1-1009 (TTAGTAGGCGAACTTACGTCGTTATGTTGGCGTTTTTCTGCACTGT), HR4X-NR4A1-1029, (CTTGTTAGCCAGGCAGATGTACTTATACGTCGAGTTGAACATAAGTGCG), HL4X-NR4A1-1486 (TTAGTAGGCGAACTTACGTCGTTATGTCGCCTGGCTTAGACCTGTA), HR4X-NR4A1-1506 (AGCAGAAGATGAGCTTGCCCTTATACGTCGAGTTGAACATAAGTGCG), HL4X-NR4A1-421 (TTAGTAGGCGAACTTACGTCGTTATGTCGAACTTGAAGGAGGCAGA), HR4X-NR4A1-441 (AGCCGTACACCTGGAAGTCCTTATACGTCGAGTTGAACATAAGTGCG) The 25 nmol H-probe oligonucleotides were ordered from IDT with standard desalting. The 100 nmol B and C oligonucleotides were ordered from IDT with HPLC purification. B and C oligonucleotides were phosphorylated with T4 polynucleotide kinase (NEB, Cat #M0201L) according to the manufacturer recommendations. Imager oligonucleotide was ordered from IDT with HPLC purification and Cy5 conjugation.

PLISH barcoding buffers and H cocktail for NR4A1 were used as previously described^30^. BC cocktail was prepared by mixing phosphorylated B and C oligonucleotides in labeling buffer (2X SSC, 20% formamide) at a final concentration of 6µM each. PLISH barcoding was performed in sealed chambers. The workflow was as follows: (1) samples were incubated in the H cocktail at 37°C for 1.5 hr, then washed 3× 5 min with H-probe buffer at RT, and then incubated in a high salt buffer (250mM NaCl, 50mM Tris pH 7.4, 2mM EDTA) at 37°C for 10 min; (2) samples were incubated in the BC cocktail + 0.2 mg/mL Heparin at 37°C for 1 hr, followed by 2 × 2 min washes with labeling buffer; (3) samples were incubated in ligation buffer at 37°C for 2 hr, followed by 2 × 2 min washes with labeling buffer; (4) sample was washed with 1X Nxgen phi29 polymerase buffer at RT for 2 min, then incubated in RCA buffer at 37°C for 2 hr, followed by 2 × 2 min washes with labeling buffer; (5) sample was incubated in labeling buffer + 0.2 mg/mL heparin + 100nM Cy5-imaging probe at 37°C for 30 min, followed by 2 × 3 min washes with PBST, and mounted in ProlongGold with DAPI (Thermo Scientific, P36931). The following day, samples were imaged on the Andor Dragonfly Spinning Disk Confocal microscope. Images were taken with 63x objective (Rapalog experiments) or 40x objective (Dyngo-4a experiments). Z stacks were taken with 1.5µm step size and max/min of the stack was determined per image by identifying the height of cells. Images were imported into FIJI and the maximum intensity projections of the channels were saved and exported as TIFF files. DAPI and Cy5 TIFF files were imported into CellProfiler^48^ and a custom pipeline was created to measure RNA signal intensities at the single-cell level. Briefly, nuclear boundaries were assigned by a shape algorithm, and then expanded by ∼10-12 micron to define sampling areas. The following data were then recorded: (i) number of nuclei and corresponding sampling areas and (ii) the coordinates of the sampling areas. For the RNA species, puncta were identified using Otsu thresholding and distinguished using an intensity algorithm. The RNA puncta were counted within determined sampling areas to calculate the number of RNA puncta per cell. For calculating % inhibition of Isoproterenol-induced response, the fold changes in samples pre-treated with Dyngo-4a versus DMSO vehicle (**Fig. 4B**) were compared to samples pre-treated with rapalog versus ethanol vehicle.

### pCRE-NLS-DD-zsGreen1 transcriptional reporter quantification

Transcriptional reporter cells were plated on Poly-L-lysine-coated coverslips and transfected with Kif1A-tdTomato-FKBP and EGFP-FRB-EEA1 using Lipofectamine 2000. When inhibiting receptor internalization, cells were first treated with 30µM Dyngo-4a or DMSO in serum-free DMEM for 20 min. When relocalizing endosomes, cells were first treated with either rapalog or ethanol for 20 min. Next, cells were then treated with Shield-1 and vehicle, isoproterenol, or isoproterenol + IBMX for 4 h. Cells were fixed with as described above and imaged on Andor Dragonfly Spinning Disk Confocal microscope. Images were imported into FIJI and individual channels were saved as TIFF files. Nuclei were identified by drawing ROIs of DAPI-stained nuclei. Cells were confirmed to have EEA1 redistribution by visual examination of the GFP channel, Kif1A-containing channel, and anti-EEA1-containing channel. In cells with efficient EEA1 redistribution, the nucleus ROI was measured for the integrated density in the GFP (reporter) channel. For calculating % inhibition of Isoproterenol-induced response, the fold changes in samples pre-treated with Dyngo-4a versus DMSO vehicle (**Fig. 4D**) were compared to samples pre-treated with rapalog versus ethanol vehicle.

### Quantitative PCR assay

For bPAC experiments, cells were transfected with each construct using the Lipofectamine 2000. On experiment day, media was changed to serum-free DMEM for 4-6 h. “Light-treated” bPACs were activated using 10% power of a 405-LED array (Sumbulbs) for indicated durations. In parallel, “no light” control samples were included. Following light activation cells, were placed in the incubator for 30 min. For Dyngo-4a experiments, HEK293 cells were incubated with DMSO or Dyngo-4a in serum-free medium for 20 min, then treated with vehicle, isoproterenol or isoproterenol/IBMX for 1 h. Total RNA was extracted from the cells using Zymo Quick-RNA MiniPrep Kit (Genesee Scientific, Cat #11-328). Reverse transcription was performed using iScript RT supermix (Biorad, Cat #1708841) following recommended manufacturer protocols. Power SYBR Green PCR MasterMix (ThermoFisher Scientific, Cat #4367659) and the following primers were used for the qPCR reactions-*GAPDH*: F 5’-CAATGACCCCTTCATTGACC-3’ and R 5’-GACAAGCTTCCCGTTCTCAG-3’; *c-fos*: F 5’-GCCTCTCTTACTACCACTCACC -3’ and R 5’-AGATGGCAGTGACCGTGGGAAT -3’; *NR4A1*: F 5’-AGTGCAGAAAAACGCCAAGT -3’ and R 5’-TTCGGACAACTTCCTTCACC -3’. Quantitative PCR was performed using CFX-384 Touch Real-Time PCR System (Biorad). All gene expression levels were normalized to the levels of housekeeping gene, *GAPDH*.

### Western Blotting

HEK293 cells were transfected with bPAC constructs using Lipofectamine 2000. The next day, cells were lysed on ice in ice-cold RIPA buffer (Sigma, Cat #R0278-500mL) with protease inhibitors cocktail (Sigma-Aldrich, Cat #P8340-5ML). Lysates were quantified by Pierce BCA Protein Assay Kit (ThermoScientific, Cat #23225), equal amounts of total protein were boiled in Laemilli dye buffer (Biorad, Cat #1610747) at 70°C for 10 min, and loaded on Mini-PROTEAN TGX Stain-Free 4-15% gels (Biorad, Cat #4568083). Gels were transferred onto nitrocellulose membranes, blocked in Tris-buffered saline (TBS)/0.05% Tween 20/5% milk, and incubated with 1:500 anti–β-actin (Santa Cruz, Cat #sc-69879) and 1:500 anti–myc (Thermo Scientific, Cat #PA1-981) in TBS/0.05% Tween 20/5% milk overnight at 4°C on a shaker. Membranes were washed, incubated with secondary antibody, 1:2500 diluted donkey anti-mouse-680 (LICOR Biosciences, Cat #926-68072) and 1:2500 diluted donkey anti-rabbit-800 (LICOR Biosciences, Cat #926-32213), in LICOR blocking buffer (LICOR Biosciences, Cat #927-40000) for 1 hr at room temperature, washed, and visualized Odyssey imager system (LICOR). Bands were quantified using ImageStudioLite. Expression of bPACs were normalized by dividing the intensity of the myc band by the intensity of the beta-actin band.

### cAMP ELISA

HEK293 cells transfected with bPAC constructs were stimulated with light or left in the dark (no light control). cAMP was measured using Cayman ELISA cAMP assay (VWR, Cat #75817-364) following manufacturer recommendations. Values were normalized to total protein concentration for each sample.

### PKA_cat_ localization and quantification

Cells on Poly-L-lysine-coated coverslips were transfected with Kif1A-tdTomato, EGFP-FRB-EEA1, TagBFP-PKAcat (a gift from Jin Zhang, UCSD), and PKARegIIB-mCherry (a gift from Roshanak Irannejad, UCSF) using Lipofectamine 2000. Cells were treated with either rapamycin or ethanol control for 20 min, then fixed as described above. Slides were imaged on the Andor Dragonfly Spinning Disk Confocal microscope. Images were imported into FIJI. ROIs were drawn around the cell periphery and nuclei identified in the Kif1A-containing channel. Clusters of EEA1-containing endosomes were segmented using the Morphological Segmentation plugin (MorphoLibJ)^46^. PKAcat particles were analyzed by thresholding the PKAcat-containing 405 channel and analyzing particles to be greater than 0.01 microns. Using the distance formula, the distance from segmented EEA1 particles and PKA participles to the nucleus center of mass was calculated based on particle and nuclear x,y coordinates.

## Author Contributions

N.G.T supervised the project. N.G.T and B.K.A.W. conceived the project and designed experiments. B.K.A.W. performed and analyzed all experiments. N.G.T and B.K.A.W. interpreted results, wrote, and edited the manuscript.

## Materials & Correspondence

Correspondence should be directed to Dr. Nikoleta G. Tsvetanova for primary data and materials.

## Acknowledgements

We would like to thank members of the Tsvetanova lab and Roshanak Irannejad (UCSF, CA) for their discussions on this project and feedback on the manuscript. Immunofluorescence microscopy imaging was performed using resources from the Duke Light Microscopy Core Facility, with specific training under Lisa Cameron, Yasheng Gao, and Benjamin Carlson. PKARegIIB-mCherry was a gift from Roshanak Irannejad (UCSF, CA). TagBFP-PKAcat was a gift from Jin Zhang (UCSD, CA). SSF-B2AR and Endo-bPAC were gifts from Mark Von Zastrow (UCSF, CA). MK1200 and dCas9-BFP-KRAB were gifts from Martin Kampmann (UCSF, CA). pBa.Kif1a 1-396.GFP was a gift from Gary Banker & Marvin Bentley (Addgene plasmid # 45058; http://n2t.net/addgene:45058; RRID:Addgene_45058). pBa-KIF5C 559-tdTomato-FKBP was a gift from Gary Banker & Marvin Bentley (Addgene plasmid # 64211; http://n2t.net/addgene:64211; RRID:Addgene_64211). pEGFP-FRB was a gift from Klaus Hahn (Addgene plasmid # 25919; http://n2t.net/addgene:25919; RRID:Addgene_25919). GFP-EEA1 wt was a gift from Silvia Corvera (Addgene plasmid # 42307; http://n2t.net/addgene:42307; RRID:Addgene_42307). Figure 1A and Figure 6E were created using BioRender. Research reported in this publication was supported by the National Institute of Health (R01NS127847 and R35GM142640 to N.G.T. and F31NS120567 to B.K.A.W.).

## Supplementary Data

**Supplementary Video 1. Live cell imaging of HEK293 cells expressing CID components**. Video shows endosome distribution after 5 min of 250nM rapalog addition.

**Supplementary Figure 1.**
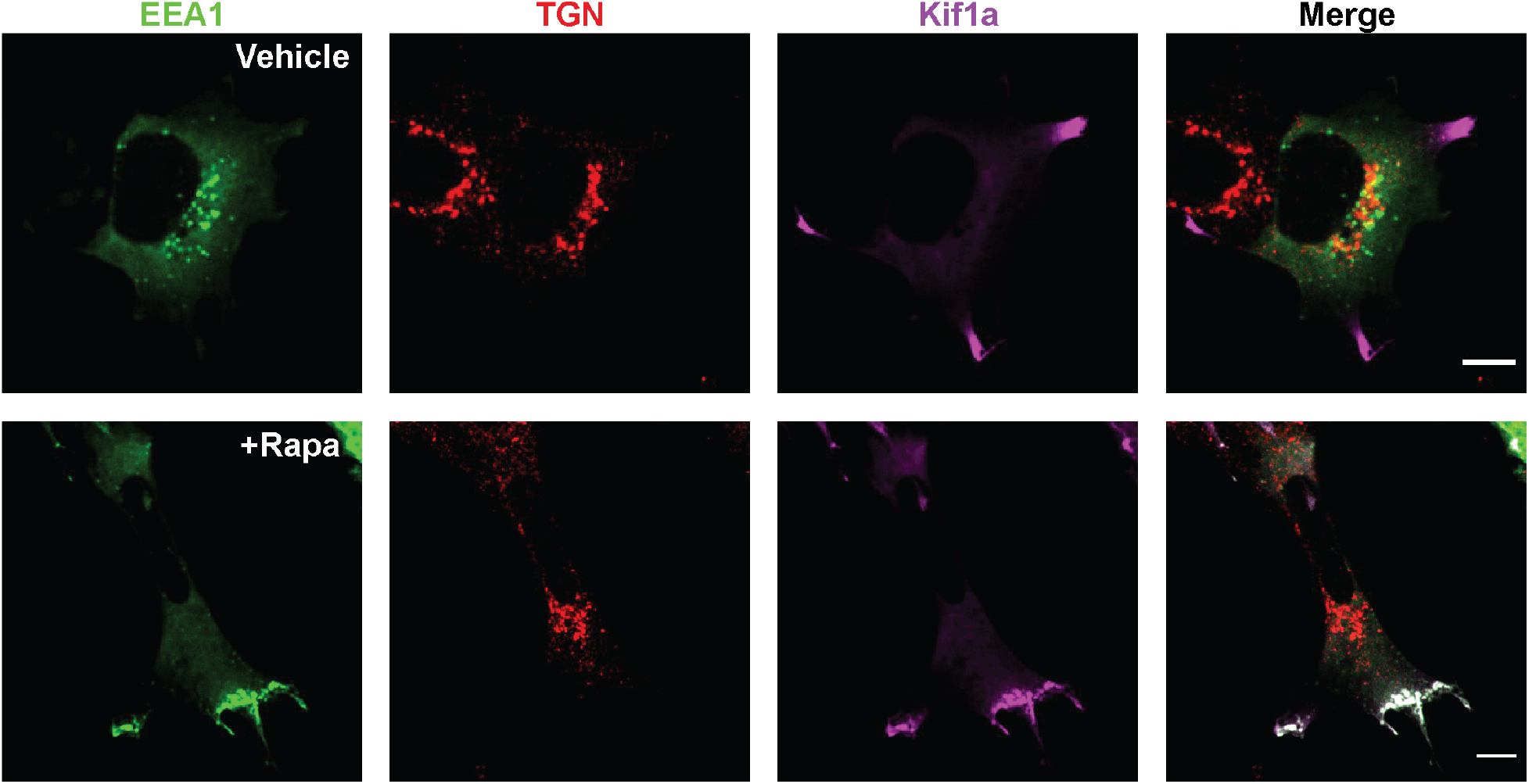
The CID approach does not impact Golgi localization. EEA1-FRB (green), localizing to early endosomes, Kif1a-FKBP (magenta), localizing near at the cell edge, and TGN marker (red) visualized by immunofluorescence microscopy in fixed cells. Cells were treated with either ethanol (“Vehicle”, top) or 1µM rapamycin (“Rapa”, bottom) for 20 min. Scale bar = 10µm.

**Supplementary Figure 2.**
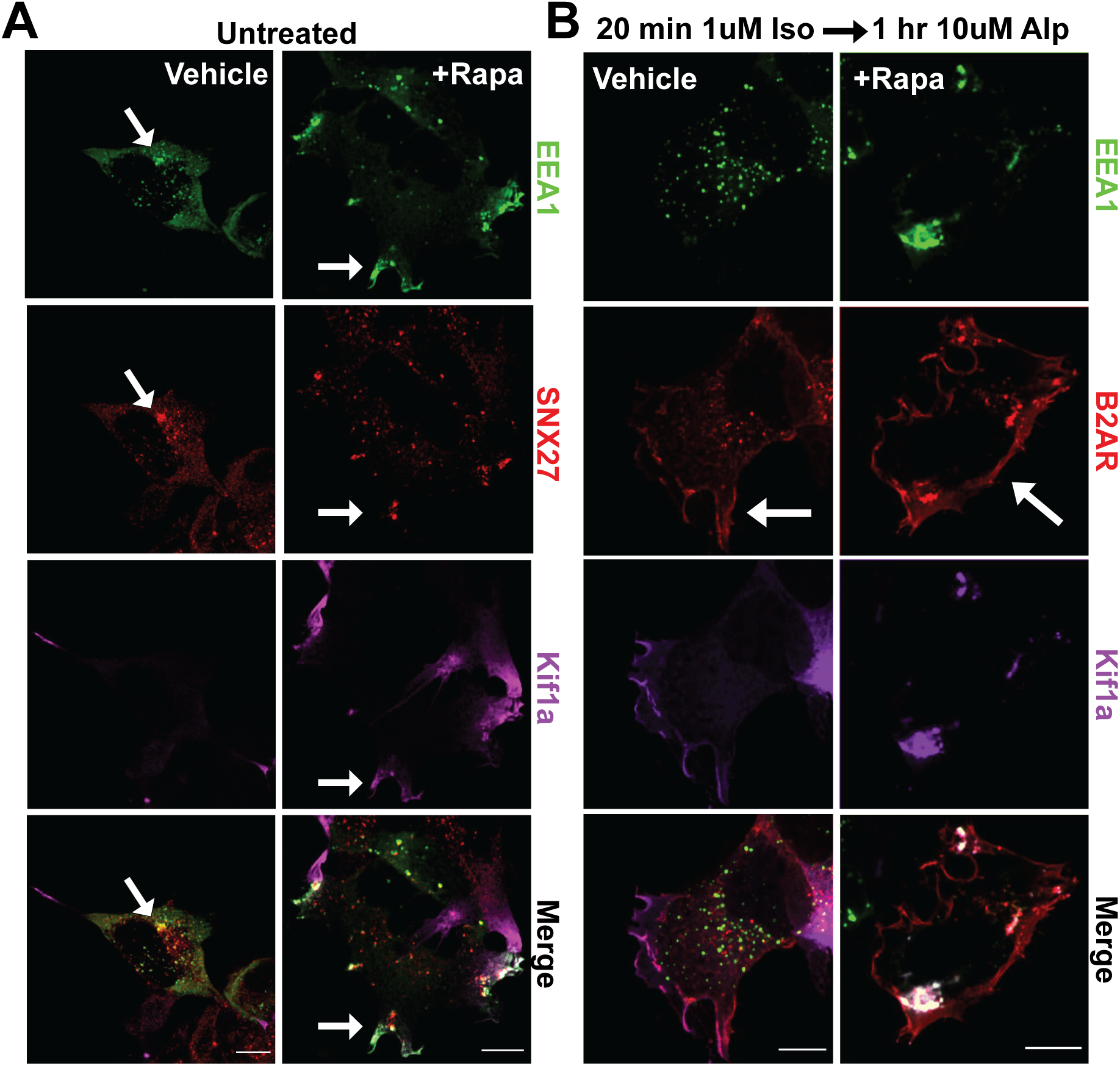
The CID approach does not alter the biochemical composition of early endosomes. **A-B**. HEK293 cells were pre-treated with ethanol (“Vehicle”) or 1µM rapamycin (“+Rapa”) for 30 min. Cells were fixed and imaged by immunofluorescence microscopy. **A**. SNX27 (red) co-localizes with early endosomes (green) in both vehicle- and rapamycin-treated cells. White arrows indicate representative co-localization between EEA1 and SNX27. **B**. Fixed-cell images of Flag-β2-AR (red) recycling after 20 min of 1µM isoproterenol-stimulated internalization into early endosomes followed by 1 hr of 10µM alprenolol treatment visualized by immunofluorescence microscopy. White arrows highlight β2-AR localization. Scale bar = 10µm.

**Supplementary Figure 3.**
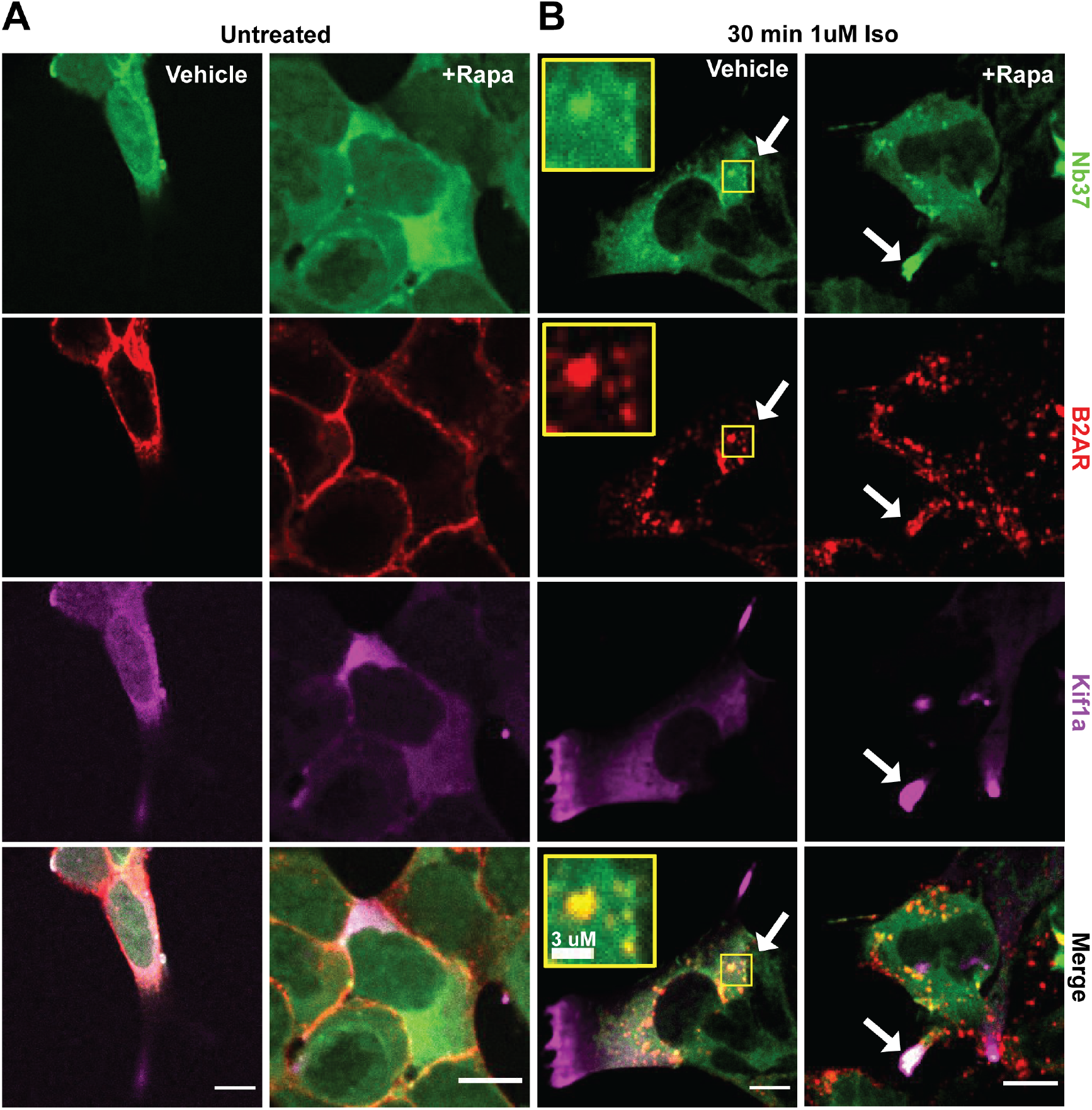
Intact β2-AR signaling in repositioned endosomes. **A-B**. HEK293 cells were pre-treated with ethanol (“Vehicle”) or 1µM rapamycin (“+Rapa”) for 30 min. Flag-β2-AR (red) and Nb37 (green) localization in untreated (**A**) and agonist-stimulated CID-expressing cells (**B**). **A**. Nb37 is cytosolic in the absence of β2-AR stimulation. **B**. Redistributed endosomes containing Flag-β2-AR colocalize with Nb37 following 1µM isoproterenol treatment for 30 min. Inset: representative co-localization region, Scale bar = 3µm. White arrows indicate examples of colocalization between Nb37 with β2-AR. Scale bar = 10µm.

**Supplementary Figure 4.**
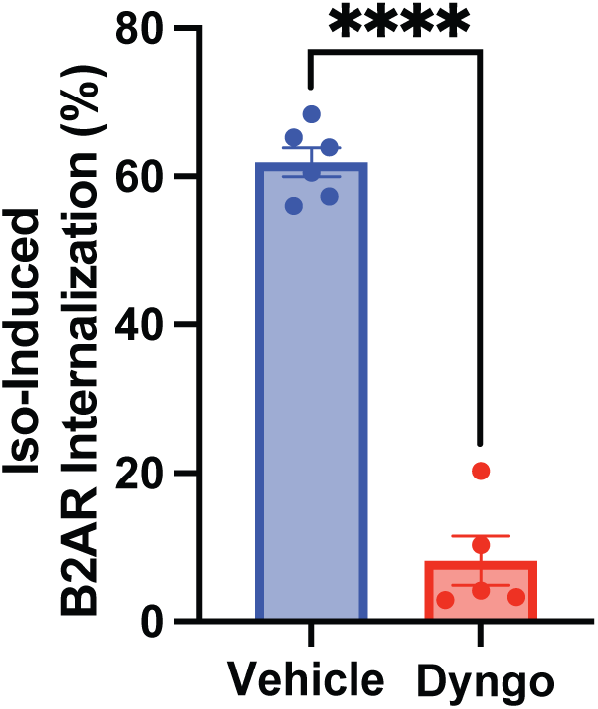
Dyngo-4a treatment inhibits agonist-Induced β2-AR internalization. HEK293 cells stably expressing Flag-β2-AR were pre-treated with DMSO (“Vehicle”) or 30µM Dyngo-4a (“Dyngo”) for 20 min before 1µM isoproterenol stimulation for 30 min. Surface β2-AR expression was measured by Alexa 647-conjugated Flag antibody and quantified by flow cytometry. Data are mean from n = 5-6 experiments ± s.e.m. **** = *p* < 0.0001 by unpaired Student’s *t-*test.

**Supplementary Figure 5.**
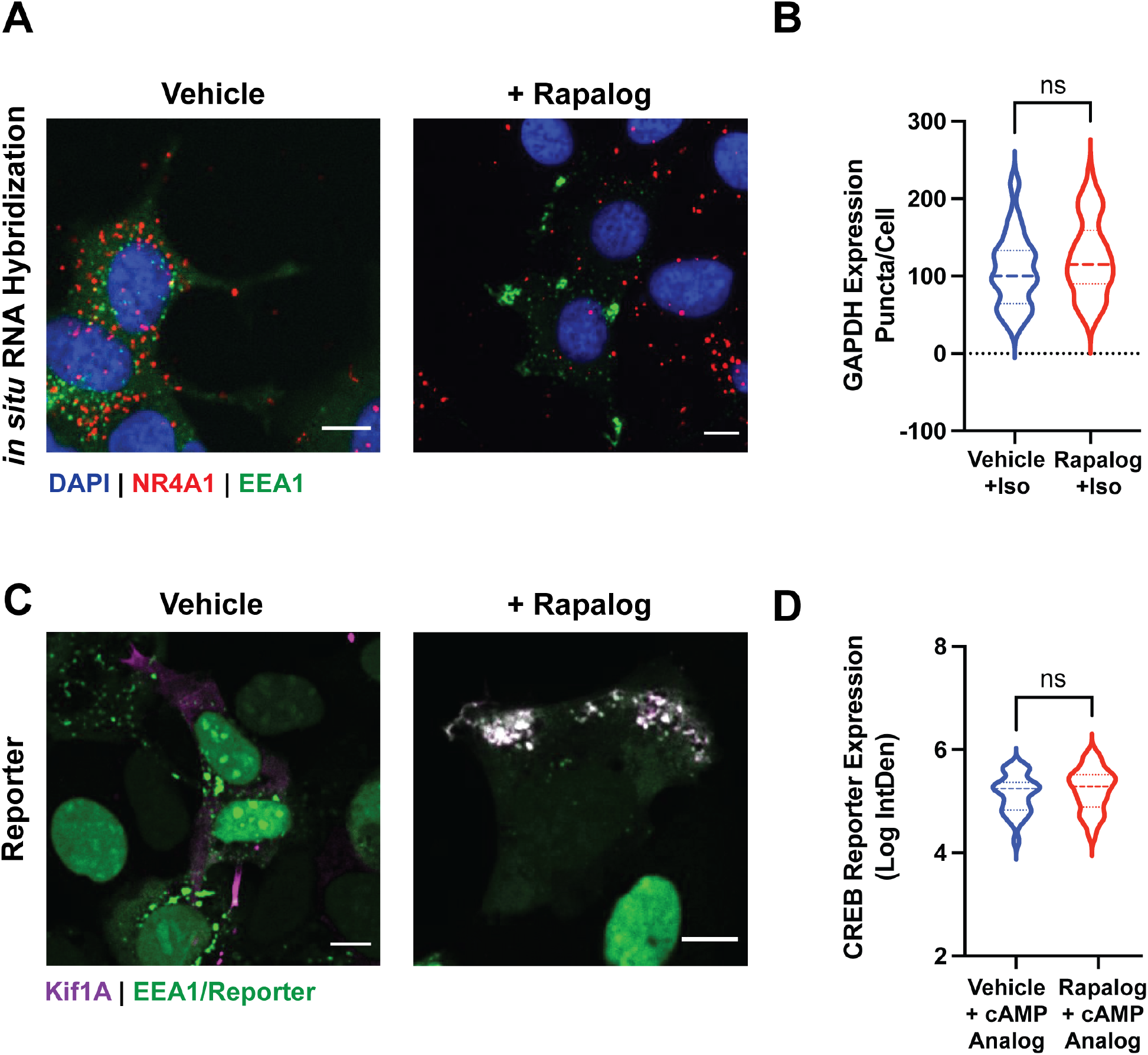
Analysis of transcriptional signaling upon endosome redistribution. **A-D**. Cells expressing the CID system were pre-treated with ethanol (“Vehicle”) or 1µM AP21967 (“Rapalog) for 30 min before agonist stimulation. **A**. Representative images of *NR4A1* expression (red puncta) upon β2-AR activation with 1µM isoproterenol for 2 hr measured by PLISH analysis. **B**. Quantification of *GAPDH* expression by PLISH analysis in cells stimulated with 1µM isoproterenol for 2 h. Data are mean from n = 3 experiments ± s.e.m.; 60 cells total/condition; ns by unpaired Student’s *t-*test. **C**. Representative images of CREB nuclear reporter expression (green) upon β2-AR activation with 1µM isoproterenol/1µM Shield for 4 hr visualized by fixed-cell immunofluorescence microscopy. **D**. Quantified CREB reporter expression following application of 150µM 8-CPT-cAMP/1µM Shield for 4 hr. Data are mean from n = 2 experiments ± s.e.m.; 27 cells total/condition; ns by unpaired Student’s *t-*test. Scale bar = 10µm.

**Supplementary Figure 6.**
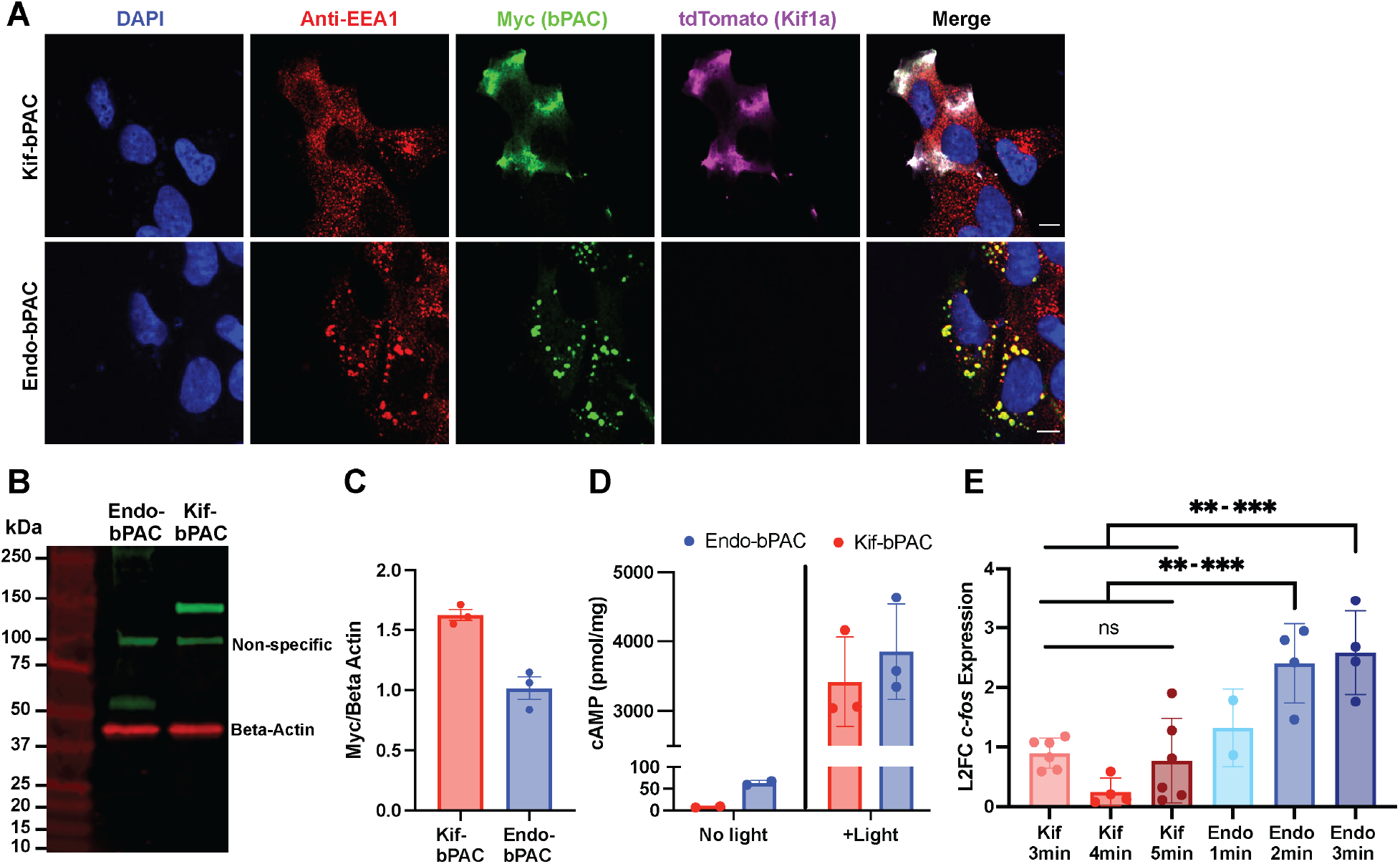
Analysis of bPAC construct localization, expression and signaling. **A**. Representative images of bPACs (green) localized to the kinesin motor (“Kif-bPAC”, top) or early endosome (“Endo-bPAC”, bottom) visualized by immunofluorescence microscopy using Alexa 647-conjugated anti-myc antibody. Relevant compartment markers (EEA1 in red, Kif1A in magenta) are shown. **B**. Representative Western blot analysis of bPAC expression (anti-myc, green) relative to beta actin control (red). A non-specific band is detected by this antibody at 100kDa. **C**. Western blot quantification. bPAC expression was normalized to beta actin levels. Data are mean from n = 3 experiments ± s.e.m. **D**. cAMP levels from Kif-bPAC and Endo-bPAC after photostimulation for 3 min of light were measured using a colorimetric immunoassay. Data were normalized to total protein concentrations. Data are mean from n = 2-3 experiments ± s.e.m. **E**. *c-Fos* induction measured by RT-qPCR analysis after bPAC photostimulation with indicated light doses. Data are mean from n = 2-6 experiments ± s.e.m. *** = *p* < 0.001, ** = *p* < 0.01 by one-way ANOVA test with Tukey. Scale bar = 10µm.

